# Deep Learning for Protein–peptide binding Prediction: Incorporating Sequence, Structural and Language Model Features

**DOI:** 10.1101/2023.09.02.556055

**Authors:** Abel Chandra, Alok Sharma, Iman Dehzangi, Tatsuhiko Tsunoda, Abdul Sattar

## Abstract

Protein-peptide interactions play a crucial role in various cellular processes and are implicated in abnormal cellular behaviors leading to diseases such as cancer. Therefore, understanding these interactions is vital for both functional genomics and drug discovery efforts. Despite a significant increase in the availability of protein-peptide complexes, experimental methods for studying these interactions remain laborious, time-consuming, and expensive. Computational methods offer a complementary approach but often fall short in terms of prediction accuracy. To address these challenges, we introduce PepCNN, a deep learning-based prediction model that incorporates structural and sequence-based information from primary protein sequences. By utilizing a combination of half-sphere exposure, position specific scoring matrices, and pre-trained transformer language model, PepCNN outperforms state-of-the-art methods in terms of specificity, precision, and AUC. The PepCNN software and datasets are publicly available at https://github.com/abelavit/PepCNN.git.

## Introduction

Protein-peptide interactions are pivotal for a myriad of cellular functions including metabolism, gene expression, and DNA replication^1,2^. These interactions are essential to cellular health but can also be implicated in pathological conditions like viral infections and cancer^3^. Understanding these interactions at a molecular level holds the potential for breakthroughs in therapeutic interventions and diagnostic methods. Remarkably, small peptides mediate approximately 40% of these crucial interactions^4^.

Traditional experimental approaches to study protein-peptide interactions, despite advances in structural biology, have significant limitations^5^. They are often costly, time-consuming, and technically challenging due to factors such as small peptide sizes^6^, weak binding affinities^7^, and peptide flexibility^8^. On the other hand, computational methods offer a complementary approach but are also encumbered by issues related to prediction accuracy and computational efficiency. This is often due to the limitations of current algorithms for the inherently complex nature of protein-peptide interactions.

Computational methods aimed at predicting protein-peptide interactions primarily belong to two distinct categories: structure-based and sequence-based. In the realm of structure-based models like PepSite^9^, SPRINT-Str^10^, and Peptimap^11^ leverage an array of structural attributes, such as Accessible Surface Area (ASA), Secondary Structure (SS), and Half-Sphere Exposure (HSE), to make their predictions. Conversely, sequence-based methods like SPRINT-Seq^12^, PepBind^13^, Visual^14^, PepNN-Seq^15^, and PepBCL^16^, utilize machine learning algorithms and various features, including amino acid sequences, physicochemical properties, and evolutionary information. Notably, PepBind^13^ was the first to incorporate intrinsic disorder into feature design, acknowledging its relevance to protein-peptide interactions^17^.

The rise of deep learning technologies has added another dimension to the computational proteomics landscape. Various algorithms now facilitate the conversion of protein features into image-like formats, making them compatible with deep learning architectures such as Convolutional Neural Network (CNN)^18^. Transformer-based models have also emerged as powerful tools for sequence representation^19^, often outperforming traditional models by capturing long-range interactions within the sequence.

For example, Wardah et al.^14^ introduced a CNN-based method called Visual, which encodes protein sequences as image-like representations to predict peptide-binding residues in proteins. Abdin et al.^15^ unveiled PepNN-Seq, a method leveraging the capabilities of a pre-trained contextualized language model named ProtBert^19^ for protein sequence embedding. Most recently, Wang et al.^16^ used ProtBert^19^ in a contrastive learning framework for predicting protein-peptide binding residues.

Deep learning algorithms, a specialized subset of machine learning, have shown considerable promise in addressing complex challenges in protein science and structural biology^20,21^. These algorithms, inspired by human cognitive processes, employ artificial neural networks to learn complex data representations^22,23^. Compared to the traditional machine learning framework like Random Forest (RF) and Support Vector Machines (SVM), deep learning models excel in autonomously discovering patterns and features from data^24^. Initially popularized in fields like medical imaging, speech recognition, computer vision, and natural language processing, these algorithms have marked milestones such as predicting folding of proteins with remarkable accuracy, making them particularly effective when applied to large and complex data^25^. Given the data-intensive nature of modern biotechnological research, proteomics is increasingly becoming a fertile ground for the application of deep learning technologies^26–28^.

CNNs^29^ have demonstrated exceptional prowess in image classification tasks, thereby suggesting their applicability to other forms of spatial data, including protein structures^30,31^. Their ability to preserve spatial hierarchies within the data makes them uniquely suited for applications in proteomics. Concurrently, advancements in natural language processing have facilitated the development of pre-trained contextualized language models specifically designed for protein biology, further enriching computational tools available for the field^32,33^.

Motivated by these technological leaps, we designed PepCNN, an innovative model that synergistically integrates protein sequence embedding from transformer language models with CNNs. Our method represents a groundbreaking, consensus-based approach by amalgamating sequence-based features derived from ProtT5-XL-UniRef50, transformer language model by Elnaggar et al.^19^ (herein called ProtT5) with traditional sequence-based (Position Specific Scoring Matrices (PSSMs)) and structure-based attributes to train a one-dimensional (1D) CNN, as shown in Figure 1. Rigorous evaluations underscore that PepCNN sets a new benchmark, outclassing existing methods such as the recent sequence-based PepBCL, PepNN-Seq that utilizes a pre-trained language model, PepBind with intrinsic disorder features, and SPRINT-Str with its emphasis on structural features like ASA, SS, and HSE. The marked superiority of PepCNN over these methodologies, in both input requirements and predictive performance, promises not only to redefine computational methods but also to accelerate drug discovery, enhance our understanding of disease mechanisms, and pioneer new computational approaches in bioinformatics.

**Figure 1.**
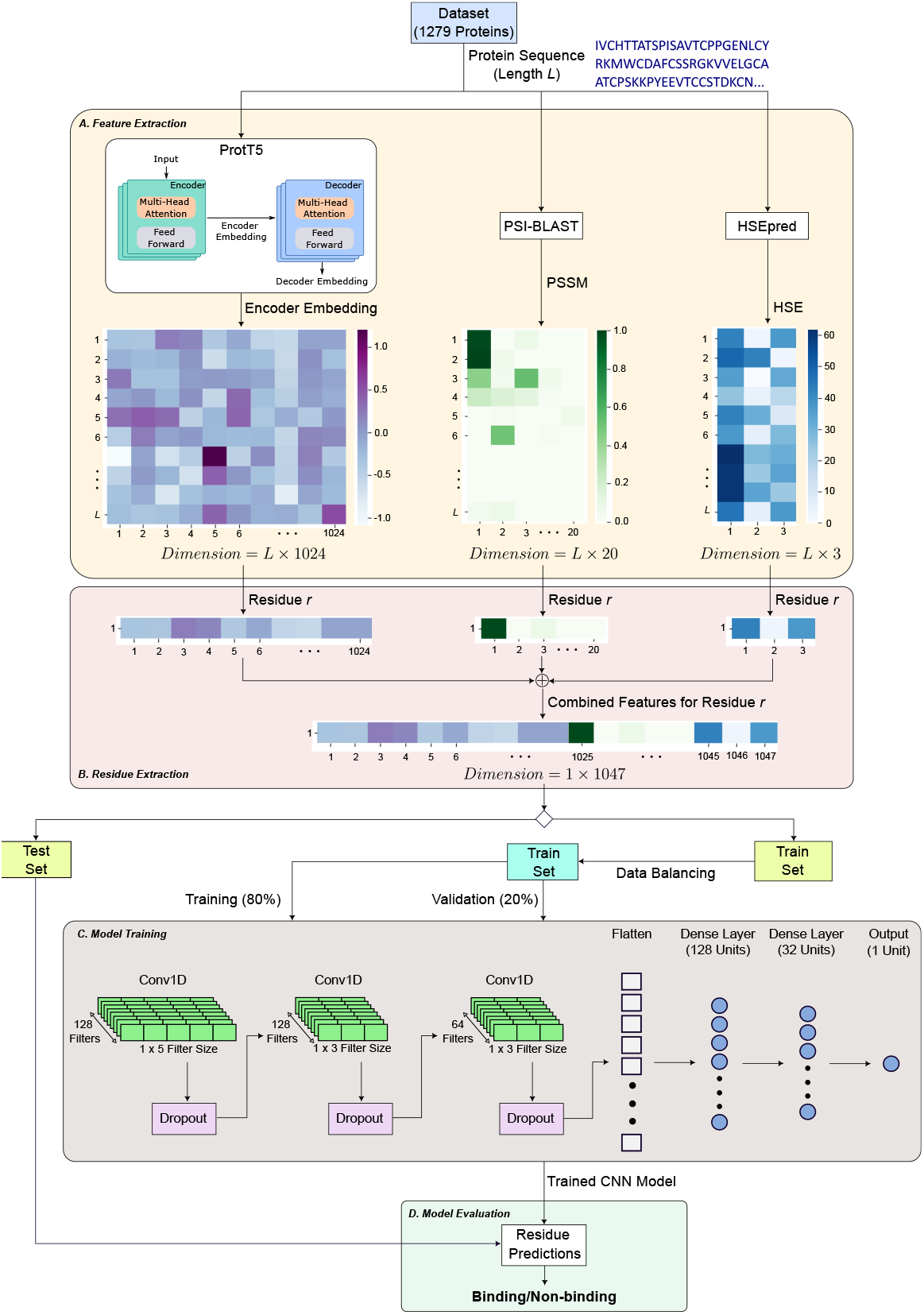
Flow diagram of the proposed work for the prediction of binding and non-binding residues. (A) Feature extraction component is where the features for each proteins are generated. (B) Residue extraction component is where the feature set pertaining to each residue is extracted. (C) The model training block contains the CNN model training step using 80% of the training set to train the network, and the remaining 20% for validation. (D) The model evaluation component is where the residues in the testing set are predicted to be binding or non-binding using the trained CNN model.

## Results

### Experimental Setup

We used two widely used benchmark datasets in this study to fairly assess and compare our proposed method with the existing approaches. These datasets are commonly used by recent state-of-the-art methods for model training and test in order to carry out evaluation and comparisons^16^. We also followed the same process for a fair comparison. The two datasets were initially obtained from the BioLiP database^34^ and sequences with a redundancy of >30% sequence identity were removed using ‘blastclust’ in the BLAST package^35^. We addressed the issue of class imbalance in our datasets by employing random under-sampling. This ensures that our model is not biased towards any particular class and can generalize well during evaluation. A residue in a protein sequence is said to be binding if any of its heavy atom is within 3.5 Å from a heavy atom in the peptide^12^ found during lab experimentation. The resulting 1,279 peptide-binding proteins contain 290,943 non-binding residues (experimental label = 0) and 16,749 binding residues (experimental label = 1). We designated the two datasets as Datasets 1 and 2, respectively, to make the discussion easier. Table 1 displays the datasets’ executive summary. The following subsections describe the specifics of the datasets for model training and evaluation.

**Table 1.**
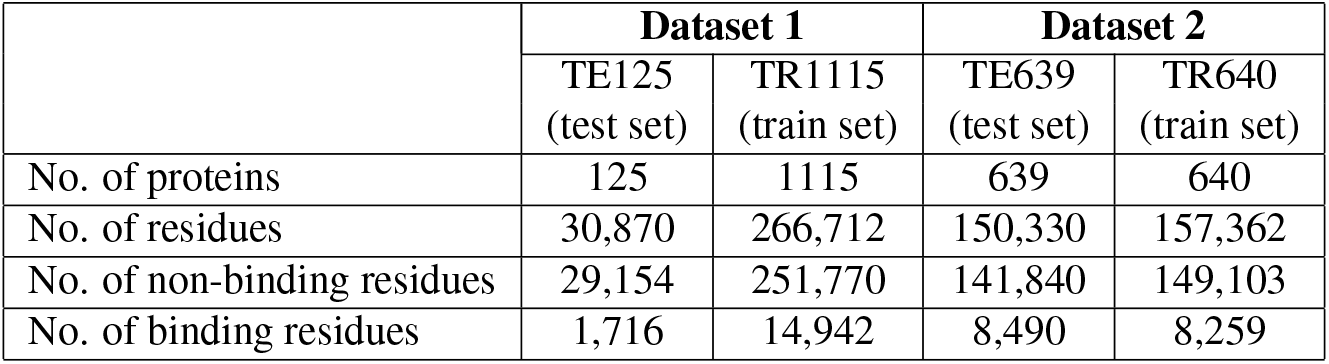
Breakdowns of Dataset 1 and Dataset 2.

|  | Dataset 1 |  | Dataset 2 |  |
| --- | --- | --- | --- | --- |
|  | TE125<br>(test set) | TR1115<br>(train set) | TE639<br>(test set) | TR640<br>(train set) |
| No. of proteins | 125 | 1115 | 639 | 640 |
| No. of residues | 30,870 | 266,712 | 150,330 | 157,362 |
| No. of non-binding residues | 29,154 | 251,770 | 141,840 | 149,103 |
| No. of binding residues | 1,716 | 14,942 | 8,490 | 8,259 |

### Dataset 1

In Dataset 1, the test set (TE125) was proposed by Taherzadeh et al.^10^ in their structure-based approach called SPRINT-Str. To create this set, they firstly selected proteins which were thirty amino acids or more in length and contained three or more binding residues. TE125 was then constructed by randomly selecting 10% of the proteins and the remaining were assigned to the training set. There are 29,154 non-binding residues and 1,716 binding residues in the 125 proteins that make up the TE125 set. In this work, we followed a similar procedure as Taherzadeh et al.^10^ to construct our training set, i.e. selecting proteins if they had more than thirty amino acids and contained three or more binding residues. As a result, 1,115 proteins were obtained for training which constituted of 251,770 non-binding residues and 14,942 binding residues. These numbers clearly show that there is an imbalance ratio of around 1:17 between the binding and non-binding residues, which can bias any model towards the classification of non-binding residues over the classification of binding residues if trained directly on this training set. Hence, the random under-sampling technique was employed to obtain the final number of non-binding residues in order to have a balanced training set. This resulted in a total of 37,355 residues for training. From this training set, 80% of the residues were actually used for training the model, and the remaining 20% of the residues were used as the validation set during the training stage.

### Dataset 2

In Dataset 2, the test set (TE639) was proposed by Zhao et al.^13^ in their sequence-based approach called PepBind. They constructed their train and test sets by randomly dividing the 1,279 proteins into two equal subsets. There were 141,840 non-binding residues and 8,490 binding residues in the 639 proteins that make up the TE639 set. In the training set, there were 640 proteins, but to save training time, 20% of the proteins were selected to train their model. The training set in this work was however created by taking all of the 640 proteins and this resulted in 149,103 non-binding residues and 8,259 binding residues. It is evident that this training set is also highly imbalanced, with an imbalance ratio of 1:18 between the binding and non-binding residues. This was also resolved by randomly under-sampling the non-binding residues in order to have a balanced training set. The final number of residues in the training set was therefore 20,647, which then underwent split with a 80:20 ratio for the final training and validation set during the model training stage.

### Comparison with Existing Methods

To show the performance of our PepCNN model, we compared the results with eight existing methods, which are: Pepsite^9^, Peptimap^11^, SPRINT-Seq^12^, SPRINT-Str^10^, PepBind^13^, Visual^14^, PepNN-Seq^15^, and PepBCL^16^. We employed sensitivity, specificity, precision, and AUC as our evaluation metrics. Sensitivity measures the true positive rate, specificity indicates the true negative rate, precision signifies the positive predictive value, and AUC represents the model’s overall classification ability. The results on TE125 and TE639 test sets are shown in tables 2 and 3, respectively. Since the test sets were also employed by these methods, their results in the tables below are taken directly from their work. As seen from the results on TE125 and TE639 test sets, PepCNN (our proposed method) achieves higher performance compared to all of the previous methods. For TE125 (Table 2) PepCNN achieves 0.254 sensitivity, 0.988 specificity, 0.55 precision, and 0.843 AUC. In comparison to all the previous methods, including the PepBCL method (the most recent and the best performing method so far), specificity, precision, and AUC have been improved by our method. The biggest improvement was seen on the AUC metric (3.4%), which is a valuable measure for the overall discriminatory capacity of the classifiers^36,37^. The results on TE639 test set is shown in Table 3 where the sensitivity, specificity, precision, and AUC values obtained by our method were 0.217, 0.986, 0.479, and 0.826, respectively. Similar results as TE125 are observed on the TE639 test set, whereby, the specificity, precision, and AUC have been increased compared to the previous methods. Again, the biggest improvement was achieved on the AUC metric (by 2.7%) compared to the previous best performing method, PepBCL. These improvements on the two test sets portray the importance of feature sets from transformer language model, PSI-BLAST (multiple sequence alignment), and structural information pertaining to half-sphere exposure and the use of this feature set with CNN to learn robust features for the prediction of binding and non-binding residues in protein sequences.

**Table 2.** Performances of the proposed PepCNN model and the previous methods on the TE125 test set. The highest values in each column are highlighted in bold.

| Methods | Sensitivity | Specificity | Precision | AUC |
| --- | --- | --- | --- | --- |
| Pepsite<sup>9</sup> | 0.180 | 0.970 | - | 0.610 |
| Peptimap<sup>11</sup> | 0.320 | 0.950 | - | 0.630 |
| SPRINT-Seq<sup>12</sup> | 0.210 | 0.960 | - | 0.680 |
| SPRINT-Str<sup>10</sup> | 0.240 | 0.980 | - | 0.780 |
| PepBind<sup>13</sup> | 0.344 | - | 0.469 | 0.793 |
| Visual<sup>14</sup> | **0.670** | 0.680 | - | 0.730 |
| PepNN-Seq<sup>15</sup> | - | - | - | 0.805 |
| PepBCL<sup>16</sup> | 0.315 | 0.984 | 0.540 | 0.815 |
| PepCNN (ours) | 0.254 | **0.988** | **0.55** | **0.843** |

**Table 3.**
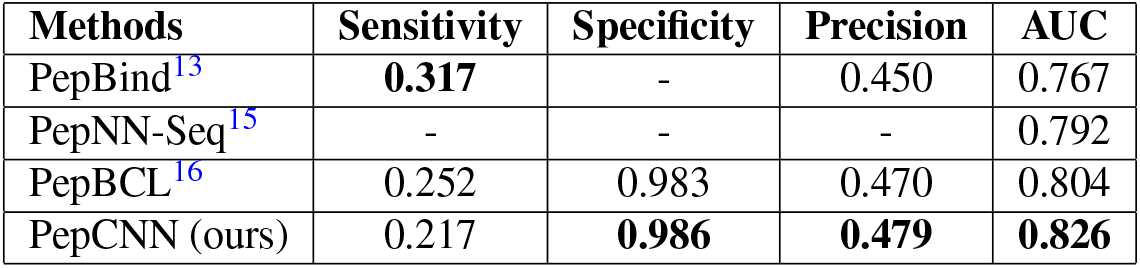
Performances of the proposed PepCNN model and the previous methods on the TE639 test set. The highest values in each column are highlighted in bold.

### Case Study

To further elaborate on the output prediction of our proposed method, we randomly selected three protein sequences from the TE125 testing set after they had been predicted by our model. These proteins were pdbID: 1dpuA, pdbID: 2bugA, and pdbID: 1uj0A and are visualized as 3D structures in Figure 2A-B, Figure 2C-D, and Figure 2E-F, respectively^38^. The *magenta* colors in the figure show the binding residues and the *gray* colors show the non-binding residues. The top visualization in the figure illustrates the experimental output (the true binding residues) of the protein, while the bottom visualization shows the binding residues of the protein predicted by our model. The protein structures B, D, and F of Figure 2 show that the predicted binding residues by our PepCNN model closely resembles the actual binding residues in the corresponding proteins (structures A, C, and E of Figure 2) detected by the lab experiment. This observation indicates a high degree of similarity between predicted and actual binding residues. This validates that our algorithm effectively leverages information from primary protein sequences for residue prediction tasks.

**Figure 2.**
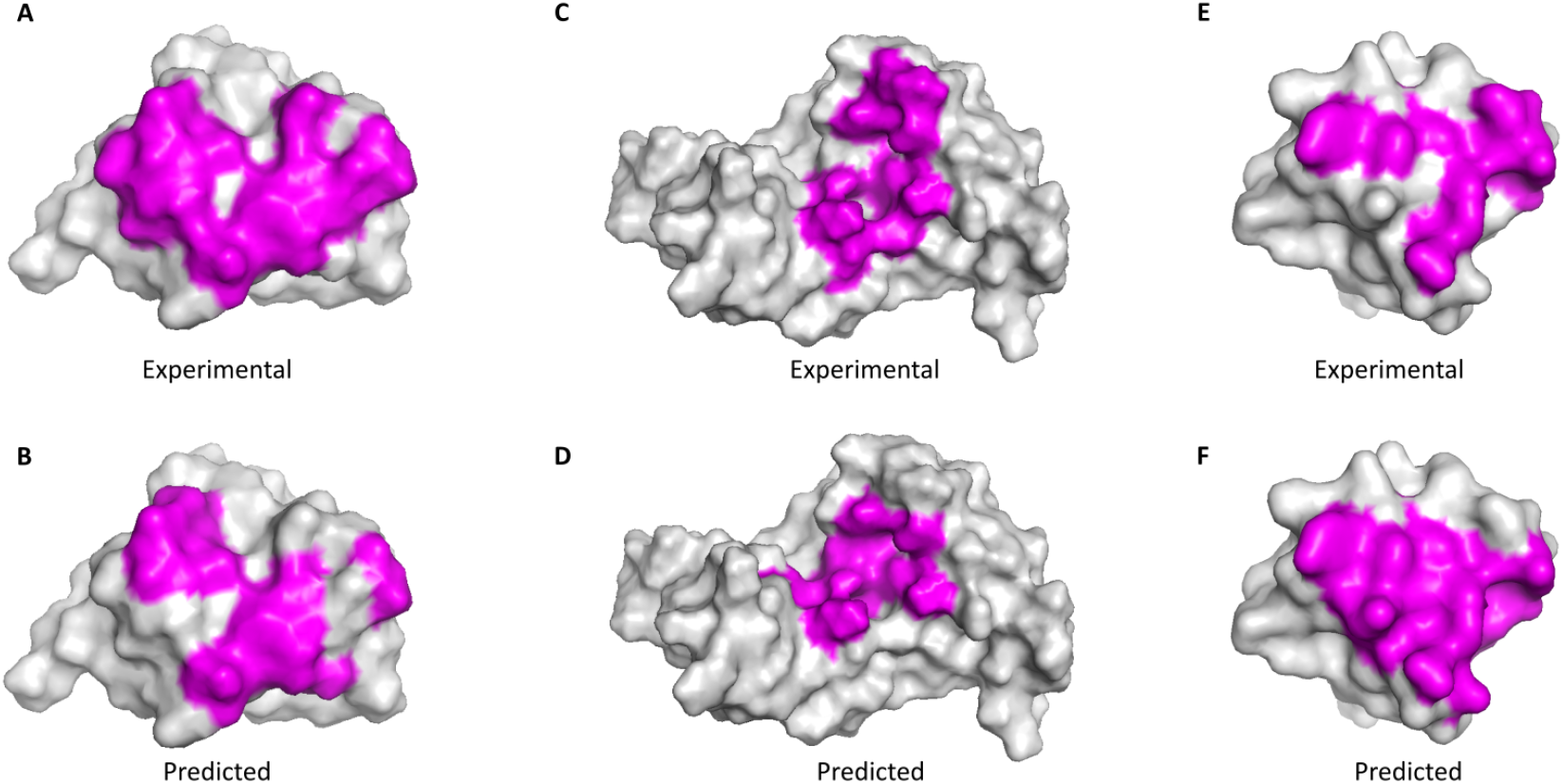
3D structure visualization of three proteins (pdbID: 1dpuA, pdbID: 2bugA, and pdbID: 1uj0A) illustrating the binding (*magenta*) and non-binding (*gray*) residues. The experimental output (true binding residues) of the proteins are located in the top part (A, C, and E) and its corresponding predicted binding residues by our method PepCNN are located in the bottom part (B, D, and F).

### Insights into the Residue Features

We had built an initial model in this work in which the performance of each of the feature sets and their combinations had been evaluated. In this initial model, we employed an ensemble of RF classifiers to have diverse training sets for Dataset 1 for a thorough evaluation and at the same time have less computational complexity compared to using a deep learning model. The ensemble consisted of 15 individual RF classifiers with different training sets by randomly selecting different non-binding residues during the data balancing stage. The hyper-parameters of the classifiers were tuned using the Hyperopt algorithm^39^ with 5-fold cross-validation scheme. The ensemble’s final predictions on the test set were to determined by averaging the individual RF classifiers’ probabilities, ensuring a robust and generalized performance.

Figure 3 shows the ROC curves obtained for the individual feature sets and the different feature set combinations on TE125. It can be seen that the embedding from the ProtT5 protein language model attains the highest AUC value out of all the individual feature sets. As the bindings are dependent on the conformations of proteins^40^, this affirms that the embedding from the pre-trained transformer model captures essential information concealed in the primary protein sequences which relates to the structure and function of proteins and therefore contributes immensely to the binding prediction. Furthermore, when the feature set combinations were assessed, it was found that the combination of Embedding, PSSM, and HSE achieved the overall best AUC value. The result obtained by combining the features suggests that PSSMs from sequence alignment and the structural properties from half-sphere exposure add more information to the protein sequence representations of the transformer model. This final feature combination was then used to build our deep learning model to further improve the performance.

**Figure 3.**
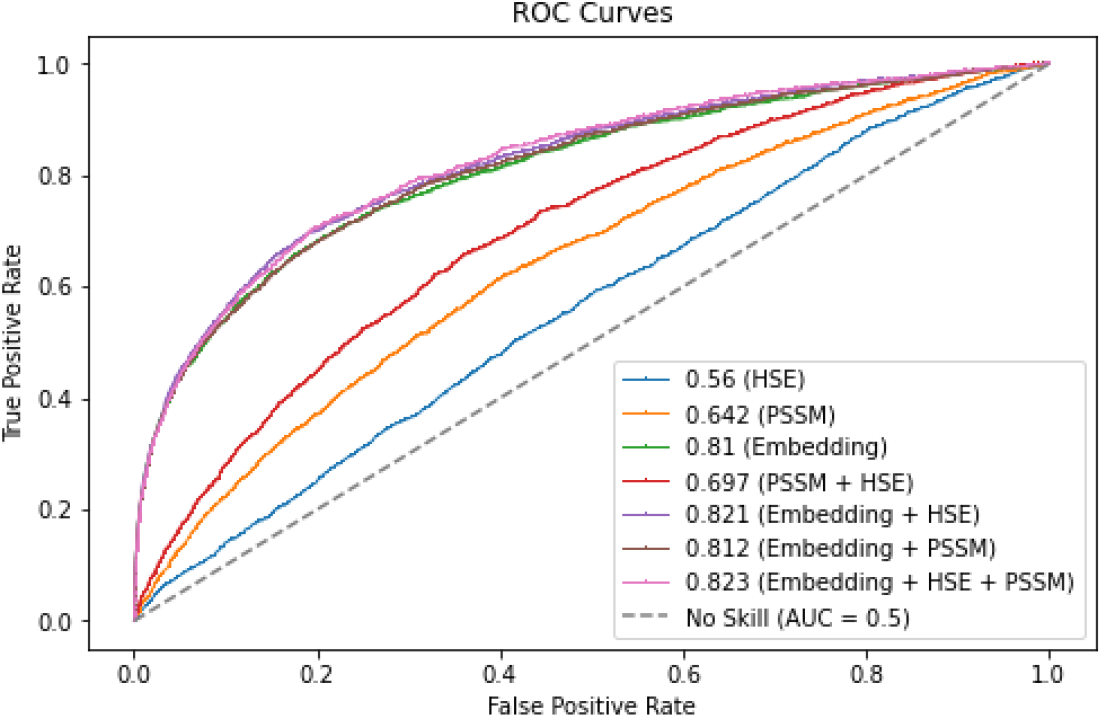
ROC curves for the individual feature sets and the different feature set combinations using the ensemble of RF classifiers on TE125.

## Discussion

We have demonstrated that PepCNN can effectively predict binding and non-binding residues in protein sequences, and thus established the possibility of the transformer embedding, PSSM, and HSE feature combination with CNN as feature extractor to predict interaction sites and explore the mechanisms of protein-peptide binding. These features enable us not only to predict interaction sites but also to explore the mechanisms underlying protein-peptide binding. The three proteins were randomly selected for structural visualization so that the similarity of the predicted and experimental binding residues could be deciphered. The strong correlation observed suggests that our approach holds promise for identifying prospective binding sites in a broad array of proteins.

When evaluating a predictor, the most ideal model would be the one which has the sensitivity and specificity measures equal to 1, however, this incidence is not prevalent in clinical and computational biology research since the measures increase when either of them decreases^41^. The ROC curve, which is an analytical method represented as a graph, is therefore mainly used for evaluating the performance of a binary classification model and to also compare the test result of two or more models. Essentially, the curve plots the coordinate points using the false positive rate (1 - specificity) as the x-axis and the true positive rate (sensitivity) as the y-axis. The closer the plot is to the upper left corner of the graph, the higher the model’s performance is since the upper left corner has sensitivity equal to 1 and the false positive rate equal to 0 (specificity is equal to 1). The desired ROC curve hence has an AUC (area under the ROC curve) equal to 1.

The study of protein-peptide binding is desired since the peptides exhibit low toxicity and posses small interface areas (as peptides are mostly 5–15 residues long^42^), making them good targets for efficacious therapeutic designs and drug discovery process^43^. In addition, peptide-like inhibitors are used for treating diabetes, cancer, and autoimmune diseases^44^. In the past, search for peptides as therapeutics was discouraged due to their short half-life and slow absorption^45^, however, these short amino acid chains are considered drug candidates once again due to the emergence of synthetic approaches which allow for changes to its biophysical and biochemical properties^46^.

Understanding the structure of protein-peptide complexes is often a prerequisite for the design of peptide-based drugs. The challenges of studying these complexes are unique compared to protein-protein and protein-ligand interactions. In protein-protein interactions, complexes are usually formed based on well-defined 3D structures, whereas in protein-ligand interactions, small ligands typically bind in deeply buried regions of proteins. Conversely, peptides often lack stable structures and usually bind with weak affinity to large, shallow pockets on protein surfaces^47^. Given these complexities, and the limitations of current experimental methods like X-ray crystallography and nuclear magnetic resonance, there is a compelling need for robust computational methods.

In summary, our work contributes to addressing these challenges by offering a highly accurate and computationally efficient method for predicting protein-peptide interaction sites. Such advances are crucial for both fundamental biological research and practical applications in drug design.

## Conclusion

In this work, we have developed a new deep learning-based protein-peptide binding residue predictor called PepCNN. The model leverages sequence-based features, which are extracted from a pre-trained transformer language model, as well as from a multiple sequence alignment tool. In addition to these, we incorporated a structure-based feature known as half-sphere exposure. Utilizing these diverse properties of protein sequences as input, our convolutional neural network was effective in learning essential features. As a result, PepCNN was able to outperform existing methods that also rely on primary protein sequence information, as demonstrated by tests on two distinct datasets.

Looking ahead, our future research aims to further enhance the model’s performance. One innovative avenue for exploration will involve integrating DeepInsight technology^18^. This technology converts feature vectors into their corresponding image representations, thus enabling the application of 2D CNN architectures. This change opens up the possibility of implementing transfer learning techniques to boost the model’s predictive power.

## Methods

### Evaluation Metrics

The proposed model in this work was evaluated using the residues in the test sets TE125 and TE639 after being trained on their respective training sets. These test sets are highly imbalanced, and for this reason, suitable metrics were chosen to effectively evaluate our model for the classification task. These metrics were Sensitivity, Specificity, and Precision. The formulation of these metrics are given below. 

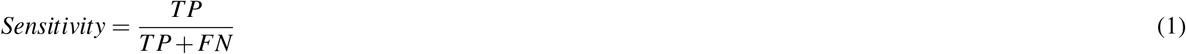

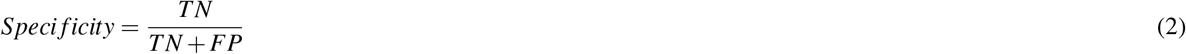

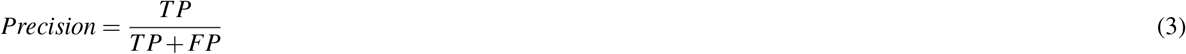

In the above formulas, TP stands for True Positives, TN stands for True Negatives, FP stands for False Positives, and FN stands for False Negatives. TP is the number of actual binding residues correctly classified by the model, TN is the number of actual non-binding residues correctly classified by the model, FP is the number of actual non-binding residues incorrectly classified by the model, and finally FN is the number of actual binding residues incorrectly classified by the model. For the given model, the Sensitivity metric (given by Eq. (1)) and the Specificity metric (given by Eq. (2)) calculate the fraction of binding residues and non-binding residues correctly predicted, respectively, and the Precision metric (given by Eq. (3)) calculates the proportion of binding residues correctly classified out of all the residues classified as binding. The values range from 0 to 1 for the metrics and the higher the value, the better the prediction model is. In addition to the above metrics, we have also included the AUC metric, which stands for the Area Under the receiver operating characteristic (ROC) Curve. AUC is a useful metric since it measures the overall performance of the classification model by calculating its separability between the predicted binding and non-binding residues. The AUC value also ranges from 0 to 1, with 0 being the worst measure of separability and 1 being a very good measure of separability.

### Feature Extraction

In the feature extraction stage of our proposed method (Figure 1A), the three different feature-types were obtained by submitting the 1,279 proteins to the three tools: ProtT5^19^, PSI-BLAST^35^, and HSEpred^48^ to acquire the Embedding, PSSM, and HSE values, respectively.

### Transformer Embedding

Among the several protein language models developed by Elnaggar et al.^19^, ProtT5 is the best performing pre-trained transformer model and is based on the T5 architecture^49^, which is akin to the originally proposed architecture for language translation task^50^ as depicted in Figure 4. It consists of the encoder and decoder blocks, where the encoder projects the input sequence to an embedding space and the decoder generates the output embedding based on the embedding of the encoder. To do this, firstly the input sequence tokens (*x*_1_, …, *x*_*n*_) are mapped by the encoder to generate representation z (*z*_1_, …, *z*_*n*_). The decoder then uses the representation z to produce output sequence (*y*_1_, …, *y*_*n*_), element by element. Both the encoder and decoder have the main components known as the multi-head attention and the feed-forward layer. The multi-head attention is a result of combining multiple self-attention modules (heads), where the self-attention is a attention mechanism that relates different positions in the input sequence to compute its representation. The attention function maps a position’s query vector and a set of key-value vectors for all the positions to an output vector. In order to carry out this operation for all the positions simultaneously, the query, key and value vectors are packed together into matrices *Q, K*, and *V*, respectively, and the output matrix is computed as: 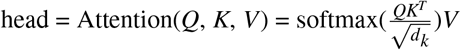 where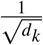 is the scaling factor. It is much beneficial to have multi-head attention instead of a single self-attention module since it allows for the capturing of information from different representations at the different positions. This is done by linearly projecting the queries, keys and values *n* times. The multi-head attention\ is therefore given by: MultiHead(*Q, K, V*) = Concat(head_1_, …, head_n_)*W*^*O*^, where 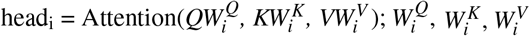 and *W*^*O*^ are projection matrices. The ProtT5 transformer used in this work is a 3 billion parameter model which was trained on the Big Fantastic Database^51^ and fine-tuned on the UniRef50^52^ database. Even though ProtT5 has both encoder and decoder blocks in its architecture, the authors found that the encoder embedding outperformed the decoder embedding on all tasks, hence the pre-trained model extracts the embedding from its encoder side. The output embedding of the ProtT5 model is a matrix of dimension *L ×*1,024 (where *L* represents the protein’s length and 1,024 the values of the network’s last hidden layer). This matrix captures relationships between amino acid residues in the input protein sequence based on the attention mechanism and produces a rich set of features that encompasses relevant protein structural and functional information.

**Figure 4.**
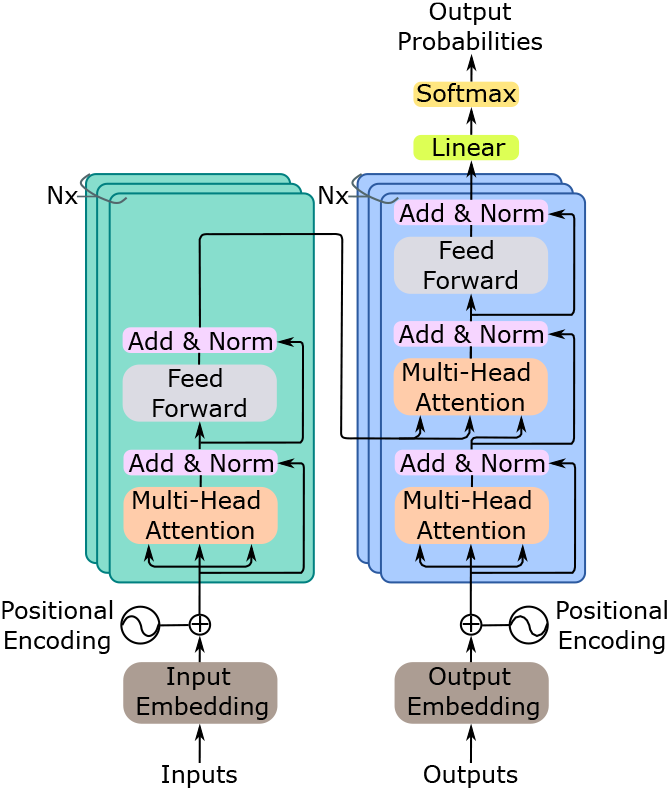
The original encoder-decoder Transformer^50^ which was proposed for language translation task. The network can have layers of these encoder-decoder modules, denoted by *Nx*. The input sequence is fed to the encoder and the decoder produces a new output sequence. At each timestep, an output is predicted, which is then fed back to the network (decoder), including all the previous outputs, to predict the output for the next timestep and so on until the output sequence (translation) is produced.

### Position Specific Scoring Matrices

PSI-BLAST is the second feature extractor method employed in this work and it was used to obtained the sequence-profiles. PSI-BLAST was run using the *E*-value threshold of 0.001 in three iterations which resulted in two matrices, log odds and linear probabilities of the amino acids, with dimensions *L×* 20 (where 20 represents the 20 different amino acids of the genetic code). The matrix with linear probabilities was used in this work in which each of the elements in the row represent the substitution probabilities of the amino acid with all the 20 amino acids in the genetic code. PSSM can therefore be formulated as *P* = {*P*_*ij*_ : *I* = 1…*L and j* = 1…20}, where *P*_*ij*_ is the probability for the *j*th amino acid in the *i*th position of the input sequence and has a high value for a highly conserved position, while a low value indicates a weakly conserved position.

### Half-Sphere Exposure

The HSE values of the proteins were obtained from the HSEpred server. This gives a measure of how buried an amino acid is in the protein’s three-dimensional structure. HSE for a residue is measured by firstly setting a sphere of radius *r*_*d*_ = 13 Å at the residue’s C*α* atom, secondly, dividing this sphere into two halves by constructing a plane perpendicular to a given C*α*-C*β* vector that goes through the residue’s C*α* atom resulting in two HSE measures: HSE-up (refers to the upper sphere in the direction of the side chain) and HSE-down (refers to the lower sphere which is in the opposite direction to the side chain), andfinally measuring the number of C*α* atoms in the upper and lower half of the sphere, respectively^48^. Refer to Figure 5 for the illustration of the HSE-up and HSE-down measures. Contact number is another important measure and it indicates the total number of C*α* atoms in the sphere of the C*α* atom of a residue^53^. The output of HSEpred is a feature matrix of dimension *L×* 3 where 3 represents to the values of HSE-up, HSE-down, and the contact number for each residue.

**Figure 5.**
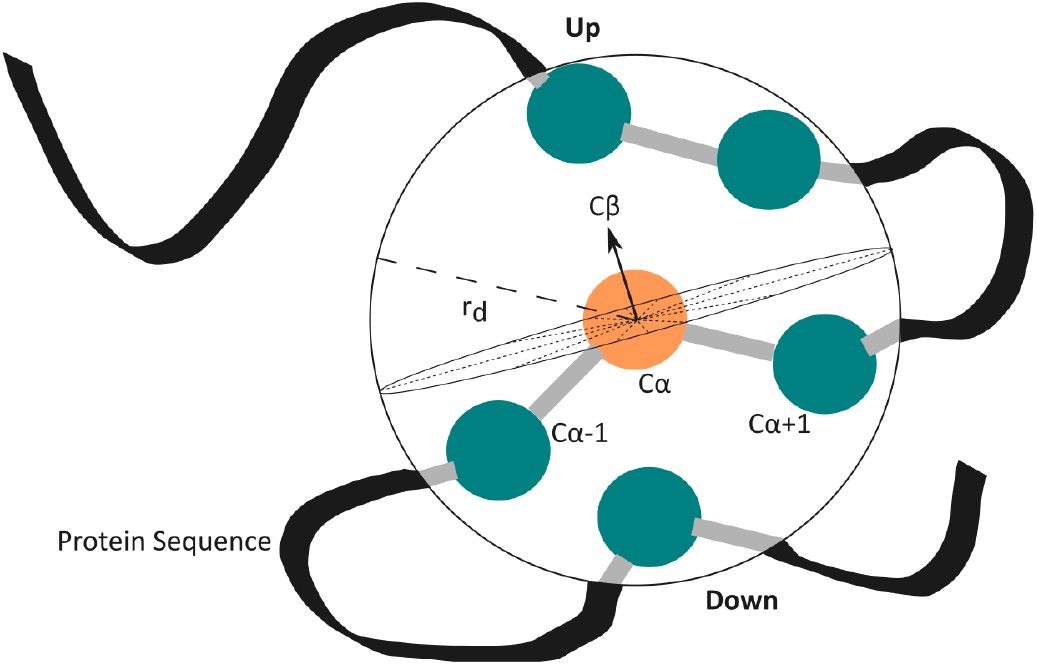
Depiction of the HSE-up and HSE-down measures. The dotted lines indicate the plane’s position which divides the sphere of the residue’s C*α* atom (*orange*) with radius *r*_*d*_ into two equal half spheres. The other C*α* atoms (*green*) represent part of other residues in the protein sequence.

### Convolutional Neural Network

From the deep learning area, CNN is one of the most widely used network in the recent times^54^. It is a type of feed-forward neural network that uses convolutional structures to extract features from data. A CNN has three main components: convolutional layer, pooling layer, and fully connected layers. The convolutional layer consists of several convolution filters. It produces what are known as feature maps by convolving the input with a filter and then applying nonlinear activation function to each of the resulting elements. The border information can be lost during the convolution process, so to mitigate this, padding is introduced to increase the input with a zero value, which can indirectly change its size. Additionally, the stride is used to control the convolving density. The density is lower for longer strides. The pooling layer down-samples an image, which reduces the amount of data and at the same time preserves useful information. Moreover, by eliminating superfluous features, it can also lower the number of model parameters. Fully connected layers are added after several convolutional and pooling layers. In the fully connected layers, all the previous layer neurons are connected to every neurons in the current layer and this results in the generation of global semantic information. The network can more accurately approximate the target function by increasing its depth, however, this also makes the network more complex, which makes it harder to optimize and are more likely to overfit. CNN has made some outstanding advancements in a variety of fields, including, but not limited to, computer vision and natural language processing, which has garnered significant interest from researchers in various fields. A CNN can also be applied to 1D and multidimensional input data in addition to the processing of 2D images. In order to process 1D data, CNN typically uses 1D convolutional filters (as portrayed in Figure 6).

**Figure 6.**
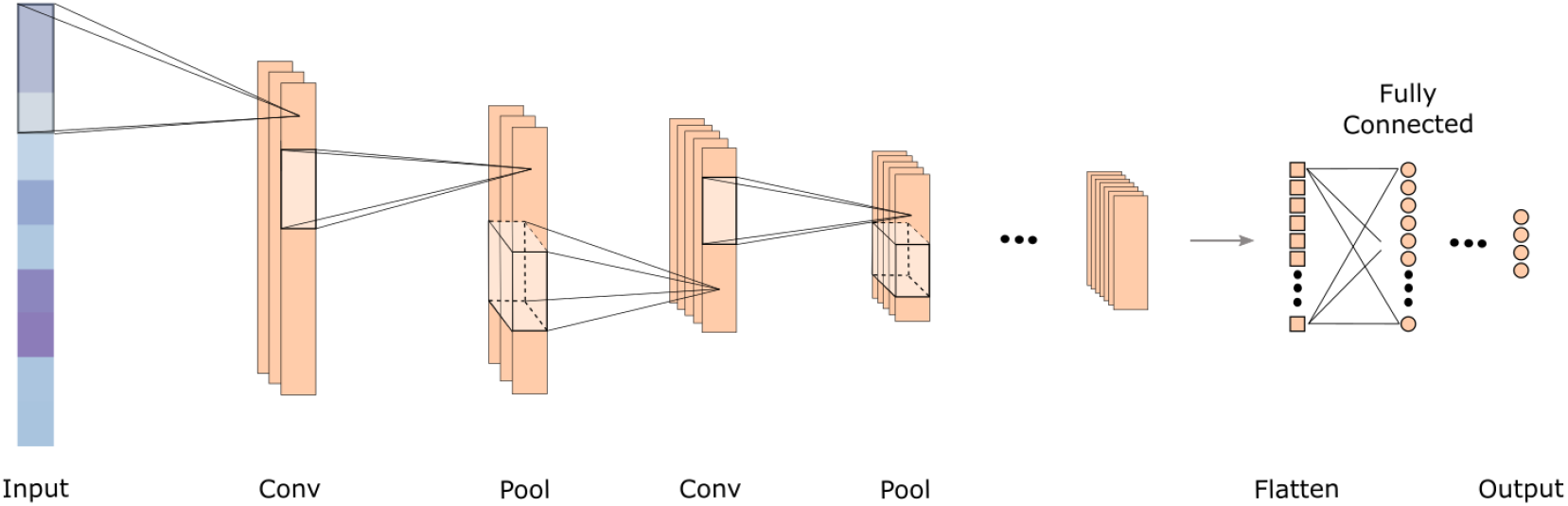
A sample 1D CNN depiction which shows the flow of information from the input to the output through its three main layers: convolutional, pooling, and fully connected.

### Building the Deep Learning Model

In order to build a classifier that carries out per residue binding/non-binding prediction, it is important to extract information pertaining to each residue. In the residue extraction stage of our proposed method (Figure 1B), we represented each residue with its sequence based (pre-trained transformer embedding and PSSM) and structure (HSE) based information. This was done by extracting the values corresponding to each residue from the three feature matrices obtained when the proteins were submitted to the three feature extraction tools. Tensor sum was applied to the resulting vectors, i.e. 1*×* 1,024 Embedding vector, 1 *×*20 PSSM vector, and 1*×* 3 HSE vector, which formed a feature vector of dimension 1 *×*1,047 to represent each residue. These residues were kept in their respective sets (i.e. train and test) to effectively train and evaluate the model without bias. In the model training stage (Figure 1C), we trained a 1D CNN to build our predictor based on the Tensorflow framework^55^. The model has 8.7 million trainable parameters which were trained using 80% of the training set, and the remaining 20% were used for network validation. The model is composed of three 1D convolutional layers and two fully connected (dense) layers. For the convolutional layers, the first layer contains 128 filters of size 5, the second layer contains 128 filters of size 3, and the third layer contains 64 filters of size 3. The stride for each layer was kept as 1 and the padding was used such that the output size of each layer was equal to the input size to the layer. Dropouts were used after each convolutional layer. In the fully connected layers, the first layer and the second layer contains 128 and 32 neurons, respectively. Finally, the output was made of a single neuron for binary classification. The ReLU activation function was used in each of the layers, while a sigmoid activation function was used in the output neuron. The model was trained using Adam optimizer with a learning rate of 1 *×*10^-6^, loss using binary crossentropy, and metric as AUC. Moreover, early stopping was employed with a patience of 3. The network was optimized using the Bayesian Optimization algorithm in the Keras Tuner library^56^.

## Author contributions statement

A.Sharma and I.D curated the data and A.C. performed analysis and experiments. A.C. and A.Sharma conceived and wrote the first manuscript. T.T. and A.Sattar contributed in manuscript write-up. All authors read and approved the final manuscript.

